# Long read sequencing reveals atypical mitochondrial genome structure in a New Zealand marine isopod

**DOI:** 10.1101/2021.09.27.462060

**Authors:** William S. Pearman, Sarah J. Wells, James Dale, Olin K. Silander, Nikki E. Freed

## Abstract

Most animal mitochondrial genomes are small, circular, and structurally conserved. However, recent work indicates that diverse taxa possess unusual mitochondrial genomes. In Isopoda, species in multiple lineages have atypical and rearranged mitochondrial genomes. However, more species of this speciose taxon need to be evaluated to understand the evolutionary origins of atypical mitochondrial genomes in this group. In this study, we report the presence of an atypical mitochondrial structure in the New Zealand endemic marine isopod, *Isocladus armatus*. Data from long and short read DNA sequencing, suggests that *I. armatus* has two mitochondrial chromosomes. The first chromosome consists of two mitochondrial genomes that have been inverted and fused together in a circular form, and the second chromosome consists of a single mitochondrial genome in a linearized form. This atypical mitochondrial structure has been detected in other isopod lineages, and our data from an additional divergent isopod lineage (Sphaeromatidae) lends support to the hypothesis that atypical structure evolved early in the evolution of Isopoda. Additionally, we find that a heteroplasmic site previously observed across many species within Isopoda is absent in *I. armatus,* but confirm the presence of two heteroplasmic sites recently reported in two other isopod species.

## Introduction

Mitochondrial genomes display a diversity of structure across Eukaryotes (reviewed in Burger et al., 2003), varying from multiple circular chromosomes to single linear chromosomes. However, within Bilateria, mitochondrial genomes tend to be circular in structure and contain 37 genes (13 protein-coding, two rRNAs, and 22 tRNAs), with a conserved arrangement (Lavrov & Pett, 2016). Here, we refer to this structure as ‘typical’. However, this structure and arrangement is not ubiquitous, as atypical mitochondrial arrangements have been found in some taxa. For example, booklice (Psocoptera) possess a multipartite mitochondrial genome consisting of two circular chromosomes (Wei et al., 2012). Thrips (Thysanoptera) also possess a multipartite mitochondrial genome with massive size asymmetry (0.9 kb and 14 kb chromosomes) (Liu et al., 2017). Tuatara, *Sphenodon punctatus,* the basally diverging lepidosaur reptile endemic to New Zealand, possesses a duplicated mitochondrial genome with a high degree of divergence between the two mitochondrial chromosomes (Macey et al., 2021).

Recent research suggests that some isopods have an atypical multipartite mitochondrial genome structure (Fig. 1) consisting of both a linear and a circular chromosome (Fig. 2) (Peccoud et al., 2017; Raimond et al., 1999). This structure is particularly common across Isopoda. The circular chromosome consists of two mitochondrial genome copies fused together in palindrome or as inverted repeats (Fig. 2A) (Peccoud et al., 2017). The second, linear, chromosome (Fig. 2B) is hypothesized to be the result of linearization and self-renaturation of a single strand of the circular chromosome during replication. Self-renaturation is possible, because the circular chromosome is made of two copies that are inverted and thus self-complementary (Peccoud et al., 2017). Aside from the presence of telomeric hairpins, this linear chromosome would be considered at a nucleotide level to be ‘typical’. Throughout this paper we will refer to the circular chromosome as the “dimer” and the linear chromosome as the “monomer”. We primarily refer to either the dimer, or the “unit” which represents the fundamental repeated unit across both mitochondrial chromosomes, alongside any unique sequence between the repeats.

**Figure 1.**
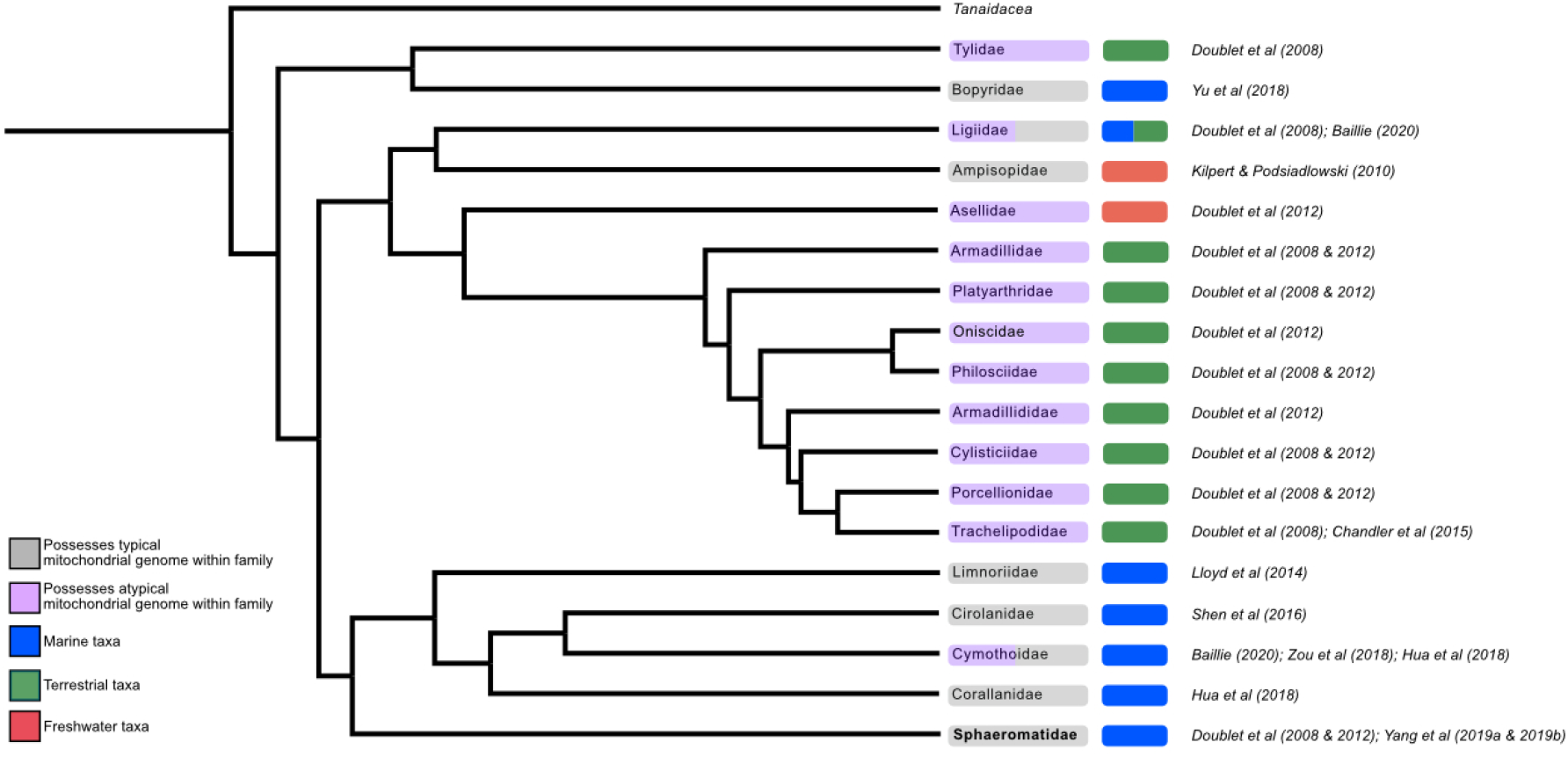
The currently known distribution of atypical mitochondrial (purple) across Isopoda at the family level. Purple indicates that for all species for which information is available, mitochondrial genomes are atypical in structure, while grey indicates they are typical. Families with purple and grey (in Cymothoidae and Ligiidae) indicate reports of both types of structure within the family. Green indicates that the taxa with the associated mitochondrial structure is terrestrial, red indicates freshwater, and blue indicates marine. This tree reflects the relationships based on Fig. S4 from (Lins et al., 2017), based on the nuclear 18S, 28S and mitochondrial COI genes. Sphaeromatidae, in bold, includes *Isocladus armatus* – which is the focus of this investigation.

**Figure 2.**
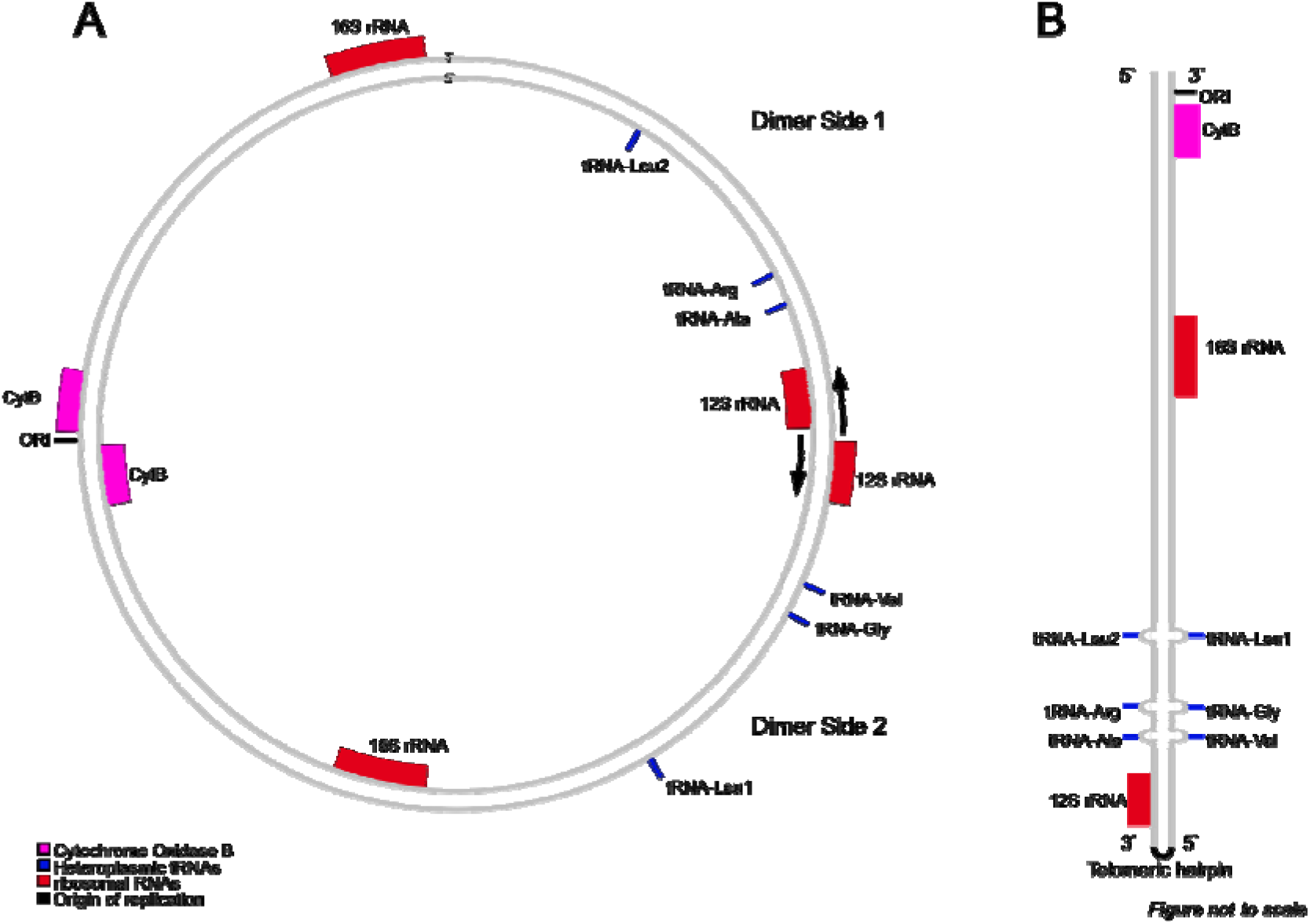
**A.** The proposed structure of the dimer. The black arrows indicate the direction of transcription, and are paired with the 12S rRNA hairpin in 2B. Putatively heteroplasmic tRNA loci are show in blue. **B.** The proposed structure of the monomer, as outlined by Peccoud et al. (2017) and Doublet et al. (2013). The monomer is a linearized copy of the dimer containing a telomeric hairpin. Importantly, there appear to be heteroplasmic sites with mismatched bases in the monomer (shown in blue, and with loops at these sites), as indicated by the presence of mirrored loci coding for different tRNAs.

These different copies or structural units of the mitochondrial genome in some isopod species are not entirely identical. Peccoud et al. (2017) have shown that there are single nucleotide differences in mirrored loci at tRNA sites, each encoding three different tRNAs. We refer to these types as either SNPs (when in reference to the mitochondrial unit) or as heteroplasmic sites (when in reference to the dimer), following the convention of Doublet et al., (2012).

This atypical mitochondrial genome is thought to have evolved prior to the divergence of suborders, such as Asellota and Oniscidea (Doublet et al., 2012). However, this hypothesis is complicated by the presence of ‘typical’ mitochondrial structure patchily dispersed in suborders such as Sphaeromatidea (family: Sphaeromatidae), Phreatoicidea (family: Amphisopodidae), and Asellota (family: Asellidae) (Fig. 1) (Doublet et al., 2012; Kilpert et al., 2012; Yang, Gao, Hui, et al., 2019; Yang, Gao, Yan, et al., 2019).

A second hypothesis has recently been proposed that relates the dominant occurrence of positive GC skews within Isopoda to the patchy distribution of mitochondrial structures across Isopoda (Baillie, 2020). This hypothesis supports the early origin of atypical structure, and proposes that an early duplication event of the mitochondrial genome resulted in an inversion of the control region (CR) and a reversal of strand skews from the ancestral negative GC skew to a positive GC skew. Occasional reversions to a ‘typical’ structure in some lineages then explain the positive GC skews seen in these species, because they necessarily only retained the single, functional, CR during the reversion event. Whilst this hypothesis necessitates regaining function of the tRNAs encoded for by heteroplasmy, there are multiple avenues for this, such as tRNA recruitment or post-transcriptional modification (Doublet et al., 2015; Sahyoun et al., 2015).

In this study, we use long read and short read DNA sequencing to investigate the structure and arrangement of the mitochondrial genome of *Isocladus armatus,* a marine isopod of the Sphaeromatidae family, which is endemic to New Zealand. We show that the *I. armatus* mitochondrial genome is atypical in structure, possessing a 28 kb circular mitochondrial genome, similar to that found in other species of isopods. Because *Isocladus armatus* is highly diverged from other known lineages to possess atypical structure, it can help to resolve the evolutionary history of isopod mitochondrial genome structure. In addition, we observe two heteroplasmic tRNA sites that have been observed previously (Chandler et al., 2015). However, we find no evidence of a third more widely studied heteroplasmy which causes a change from tRNA-Val to tRNA-Ala (Chandler et al., 2015).

## Methods

### DNA Extraction

We extracted DNA from one individual for Nanopore sequencing using a modified Qiagen DNEasy Blood and Tissue protocol, developed for *I. armatus* (Pearman et al., 2020). We extracted DNA from a second individual for Illumina sequencing using a modified Promega Wizard protocol. This protocol consisted of crushing the cephala of a specimen in a solution of chilled lysis buffer (120 μl of 0.5M EDTA and 500 μl of the provided Nuclei Lysis solution), alongside 100 μl of 1M DTT, and 30 μl of Proteinase K. The crushed sample in solution was then incubated at 65°C overnight. After overnight lysis, the sample was cooled to room temperature and 10 μl of RNAse A was added, and the sample incubated at 37°C for 30 minutes. Following this, 250 μl of protein precipitation solution was added, and the protocol was completed according to manufacturer’s instructions (page 11, Promega #TM050).

### Sequencing and Quality Control

DNA from both individuals was sequenced using both Oxford Nanopore and Illumina sequencing. For Nanopore sequencing, we followed the manufacturers protocol for native barcoding of genomic DNA for the SQK-LSK109 kit (protocol version: NBE_9065_v109_revV_14Aug2019) with a R9.4 RevD flow cell. Nanopore reads were basecalled using Guppy 3.4.3, and demultiplexed and adapters removed using PoreChop (https://github.com/rrwick/Porechop).

Illumina sequencing was carried out on an Illumina NovaSeq using 150 bp paired end reads, with an insert size of 150 bp. Potential contaminant reads from bacteria, human, or viruses were identified and discarded using Kraken2 with the maxikraken2 database (maxikraken2_1903_140GB, https://lomanlab.github.io/mockcommunity/mc_databases.html)

### Assembly

Nanopore reads were assembled into a draft genome using *Flye* (Kolmogorov et al., 2019) under default parameters with an estimated genome size of 1 GB. The mitochondrial contigs were identified by mapping all contigs to the mitochondrial genome of *Sphaeroma serratum* (Kilpert et al., 2012), resulting in the identification of a single mitochondrial contig. We then mapped all Nanopore reads to this contig, and performed a re-assembly using only the reads that mapped to the initial mitochondrial contig. For this assembly, the default settings were used with an estimated genome size of 28 kb (this size was selected based on the concordance between assembly size of the first mitochondrial contig, the size of other isopod mitochondrial genomes, and the length of the longest Nanopore reads of mitochondrial origin.).

The atypical mitochondrial structure we hypothesised (shown in Fig. 2) precluded complete polished assemblies of this mitochondrial genome, as Illumina reads were too short to be able to orient on either side of the dimer, as only reads containing the junctions can be accurately oriented. Thus, we used Geneious 9.1 (Drummond et al., 2011) to extract the ‘monomer’ from the assembly, and manually identified the primary repeat based on a self-self dotplot of the full length mitochondrial genome using YASS (Baillie, 2020; Noé & Kucherov, 2005) (Supp. Fig. 1). This was manually extracted alongside any unique sequence either side of the monomer, and this was polished three times with Illumina reads using racon (Vaser et al., 2017), and BWA (Li & Durbin, 2009) as a mapper. This contig was then visualized in IGV (Robinson et al., 2011) and three putative single base pair indels (insertions/deletions) were removed based on having coverage of less than 5% of the adjacent sites.

### Annotation

This monomer was then annotated using MITOS2 (Bernt et al., 2013), with the Al Arab protein prediction method (Al Arab et al., 2017), a non-circular assembly, a final maximum overlap of 150 bp (the number of bases that genes of different types [i.e. tRNA and rRNA] can overlap), and fragment overlap of 40 % (the fraction of the shorter sequence that can overlap with the larger sequence). These overlap values were based on existing research indicating high levels of gene overlap within isopod mitochondrial genomes (Doublet et al., 2015; Zou et al., 2018). MITOS2 also highlights potential gene duplicates and the modification of the overlap settings may increase the likelihood of erroneously identifying gene duplication. Thus, we removed annotations for potential duplicates where there was an order of magnitude difference in quality value (analogous to a BLAST e-value, low quality values are frequently spurious [http://mitos.bioinf.uni-leipzig.de/help.py]) between annotations of potential duplicates, retaining the annotation with the highest quality factor. GC-skew was calculated to confirm the position of the origin of replication using GenSkew (https://genskew.csb.univie.ac.at/), using a step size of 20 bp and window size of 100 bp (Doublet et al., 2013).

Heteroplasmic sites were identified by treating them as SNPs (Peccoud et al., 2017). These variants were identified using BWA (Li & Durbin, 2009) and bcftools (Li, 2011) with a minimum depth of 1000 X and a maximum depth of 8000 X.

## Results

### Assemblies

Nanopore sequencing on the genomic DNA from one individual was performed, and after assembly with *Flye* we identified a single mitochondrial contig. 1,817 of the Nanopore reads could be mapped back to this contig (median length 1405 bp, max length 27,851 bp). The distribution of read lengths was roughly trimodal, with peaks at approximately 1400 bp, 14,000 bp, and 28,000 bp. The peak at 1400 bp is likely the result of shearing and incorporating of low molecular weight DNA in the library, while the smaller peaks observed at 14 kb and 28 kb may result from the presence of two full length chromosomes (Supp. Fig. 2). Using *Flye* we performed a re-assembly using the 1,817 reads, resulting in a single 28,745 bp circular contig (Fig. 3), with a mean coverage of 194 X.

**Figure 3.**
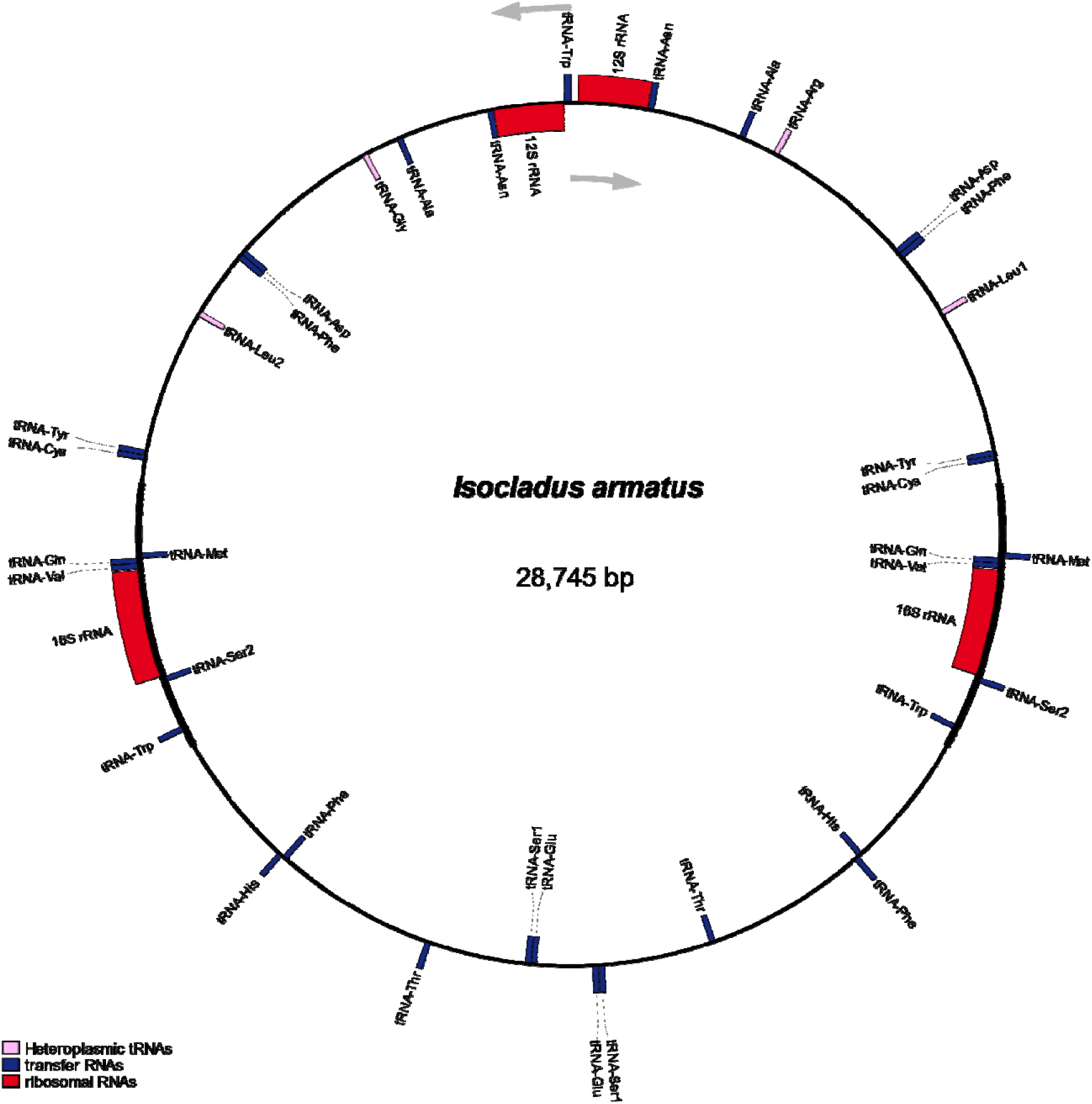
Nanopore read only assembled mitochondrial genome for *Isocladus armatus*. Grey arrows indicate the direction of transcription, blue sites indicate transfer tRNAs, pink sites indicate tRNA loci whose function varies depending on side of the dimer (heteroplasmic sites), while red sites indicate rRNAs. Annotations were created using MITOS2, and figure created using OGDRAW2.

The assembled mitochondrial genome for *Isocladus armatus* thus consists of a 28.7 kb circular chromosome consisting of two inverted repeats of a ‘typical’ mitochondrial genome. The mitochondrial unit is 14,382 bp long, and the junctions between copies of the unit comprises a total of 186 bp (Supp. Fig. 2). These junctions are located between each copy of the 12S gene, and each copy of the tRNA-Glu gene. The junction between copies of the 12S gene is 155 bp in length, while the junction between the tRNA-Glu loci is 591 bp in length.

Each unit is composed of the 13 protein-coding genes, 18 tRNAs, and 2 rRNAs. Of these tRNAs, two contain variants at mirrored sides of the dimer that enable coding for a different tRNA. Finally, another tRNA locus is found within the 16S junction, and is not duplicated. As a result, the complete dimer contains 13 duplicated protein-coding genes, 16 duplicated tRNAs, 5unique tRNAs, and 2 duplicated rRNAs (Fig. 3).

The unique component of the dimer (the ‘unit’ of the mitochondria, together with the junctions between the two copies on the dimer) was extracted, and polished with racon using Illumina data, producing a linear unit 14,569 bp long (Supp Fig. 3, GenBank: OK245257). This unit represents the repeated mitochondrial unit, alongside the unique sequences found in the junctions between repeats, and has a positive GC skew. We selected this unit as it represents the complete unique mitochondrial genome for this species, with the exception of any single base pair heteroplasmies.

The mitochondrial unit had a mean coverage with Illumina reads of 6537x (Fig. 4), containing 13 protein coding genes, 19 tRNAs, and 2 rRNAs. Two of the tRNAs appeared heteroplasmic in nature.

**Figure 4.**
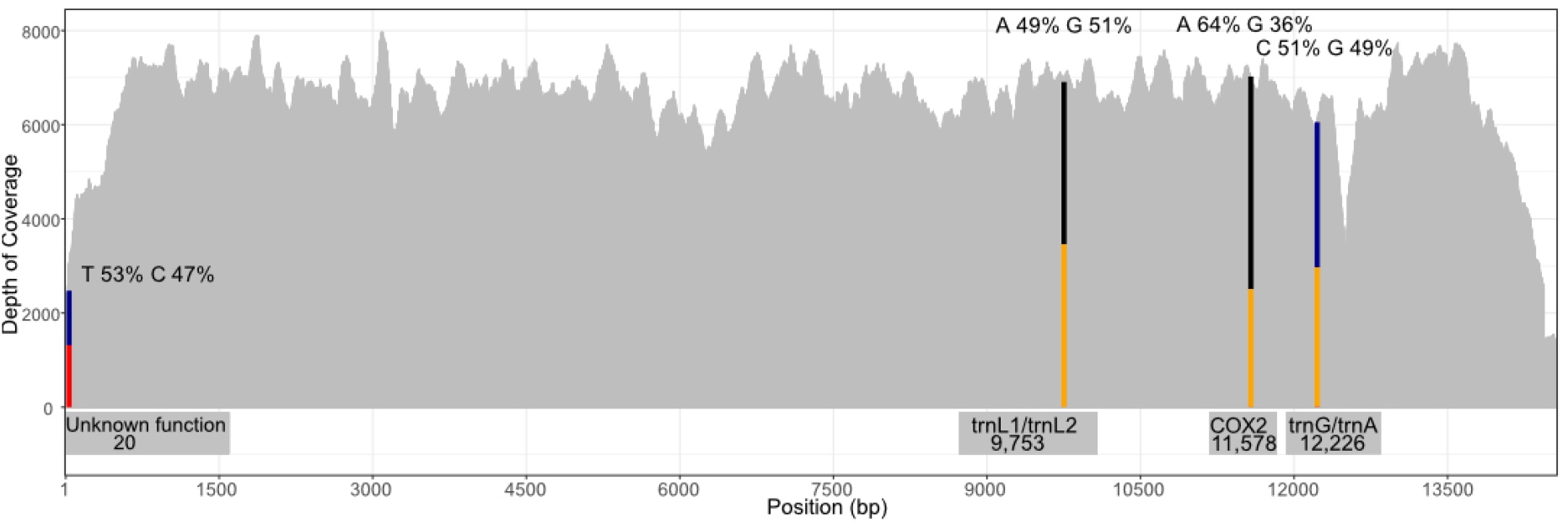
Illumina coverage of monomer, coloured bars indicate relative frequency of each base at the four SNP sites. Width of bars in these cases are not to scale. Captions within figure indicate function and position within the monomer.

### Structure of the mitogenome

We identified four single base pair mitochondrial variants (Table 1, Fig. 5) using the Illumina reads and bcftools (see Methods) (Fig. 4). Two of these SNPs were at tRNA loci located either side of the dimer, resulting in a change in the tRNA encoded at that locus. A third variant was present at an approximately 50:50 ratio within the junction between sides of the dimer. This locus has no known function. Finally, the fourth variant was a synonymous substitution in the COX3 gene, present at a 2:1 ratio. This latter variant was also conspicuously absent in the Nanopore data.

**Table 1.** ‘SNPs’ identified in the monomer using bcftools. Four SNPs have been identified, three of which appear to be present in coding regions of the mitochondria. * indicates presence only in Illumina sequencing data.

| Position | Bases | Function | Frequency of variant (%) |
| --- | --- | --- | --- |
| 20 | T-C | Non-coding | 1315:1158 (53:47) |
| 9,753 | A-G | tRNA-Leu1/tRNA-Leu2 | 3347:3460 (49:51) |
| 11,578* | A-G | Unknown CAA-CAG<br>Gln-Gln anticodon substitution in the<br>COX3 gene | 4517:2504 (64:36) |
| 12,226 | C-G | tRNA-Gly/tRNA-Arg | 3094:2963 (51:49) |

Overall, mitochondrial gene order in *I. armatus* was relatively consistent with other isopod species. As has been documented in other species of isopods, we found that the trnI locus was absent in *I. armatus* (Kilpert et al., 2012; Zou et al., 2018).

## Discussion

Using a combination of long and short read DNA sequencing, we have shown that the marine isopod, *Isocladus armatus,* exhibits an atypical mitochondrial genome structure. This atypical structure consists of a circular 28.7 kb chromosome containing two copies of a ‘typical’ mitochondria fused together as inverted repeats. This circular structure is similar in size to the mitochondrial genomes found in other isopods with atypical structure (Chandler et al., 2015; Raimond et al., 1999). We identified three sites that represent differences between copies of these repeats (termed heteroplasmies). The first site is a novel single base pair substitution that occurs in a non-coding region near the junction of the repeats and appears to be unique to *I. armatus*. The other two heteroplasmies have been previously identified in other species of isopods (Chandler et al., 2015; Peccoud et al., 2017). These two sites (positions 9,753, and 12,226) are responsible for the change in tRNA function at these loci between copies of the dimer due to a single base change in the anticodon. We also provide evidence for a putative second chromosome, which is approximately 14 kb and is, in other isopods, a linearized copy of a single mitochondrial genome (non-duplicated). We term this chromosome the ‘monomer’. While sequencing reads could not be assigned to the specific mitochondrial chromosome, read length distributions from nanopore sequence suggest the possible presence of this monomer.

The two heteroplasmic sites (tRNA-Gly-Arg, and tRNA-Leu1-Leu2) we observe were recently described in three species of Oniscid isopods (Chandler et al., 2015). The presence of these sites in *I. armatus* indicates that both the atypical structure, and heteroplasmic sites have been maintained over evolutionary time for hundreds of millions of years, as the most recent common ancestor between a Sphaeromatid and Oniscid isopod existed approximately 400 million years ago (Lins et al., 2012). Our data indicate that the heteroplasmic sites observed in *I. armatus,* have been conserved for approximately 150 million years longer than previous estimates (Chandler et al., 2015; Doublet et al., 2012). This is not particularly surprising as it is likely that this structure is maintained by balancing selection, as the loss of a tRNA would likely be lethal (Peccoud et al., 2017).

We identified heteroplasmies located on the tRNA-Arg/tRNA-Gly genes of the dimer. These sites are commonly ‘heteroplasmic’ in species with an ‘atypical’ mitochondrial genome, therefore enabling the expression of both tRNAs (Chandler et al., 2015). However, the tRNA-Arg gene is missing in some isopod species with a ‘typical’ mitochondrial genome structure (Kilpert et al., 2012; Yang, Gao, Hui, et al., 2019), while the tRNA-Gly gene is lacking in the Sphaeromatid isopod, *Sphaeroma terebrans* (M. Yang, Gao, Yan, et al., 2019). The lack of these specific tRNA loci in some isopods with ‘typical’ mitochondrial genome structures is puzzling because these lineages appear to have lost functionality of at least one tRNA locus. Despite being absent from mitochondrial gene annotations, the critical role that these tRNAs play in cellular function precludes the possibility that they are unexpressed. Instead, mechanisms such as multiple mitochondrial haplotypes or post-transcriptional modification may play a role in preserving the function of these genes. For example, Doublet et al. (2013) found evidence for multiple mitochondrial haplotypes that could ensure continued functioning of these genes in the genus *Armadillidium*. Additionally, post-transcriptional modification has been proposed as an explanation for continued functionality of other tRNAs in *Ligia oceanica* (Kilpert et al., 2012). This has also been directly observed in other isopods such as *Armadillidium vulgare,* where it has been found to influence expression of the tRNA-His locus (Doublet et al., 2015).

In addition to heteroplasmies associated with tRNA function, we observed a fourth variable site that we were unable to identify as heteroplasmic. This site was only present in the Illumina sequencing data and was conspicuously absent in the Nanopore data. As a result, we are unable to identify whether this site is variable between sides of the dimer, as short-read Illumina data would not facilitate the determination of read orientation. However, we expect the variation at this locus does not occur on the dimer (i.e. only one haplotype of the SNP is present on the dimer), because when treating these variable sites as SNPs, heteroplasmic loci appear at a 1:1 ratio, while this SNP occurs at a 2:1 ratio (with near 1:1 abundances for each SNP on the forward and reverse strands). This SNP occurs in the COX3 gene and relates to a synonymous substitution in a tRNA-Gln anticodon. We are unable to determine whether this variant is found universally in *I. armatus,* because we only sequenced two individuals, using two different sequencing approaches.

In *I. armatus,* there are two potential explanations for the presence of this SNP. Given the occurrence of this SNP at a 2:1 ratio, one possibility is that this SNP occurs on the monomer where the monomer represents c. 33% of copies of the mitochondrial unit. Peccoud et al (2017) proposed that the monomer may simply be the result of self-renaturation of the dimer during replication. Thus it would be non-functional due to the presence of mismatching bases at heteroplasmic sites on reads originating from the monomer. Alternatively, the absence of this SNP in Nanopore data, despite the presence of three other SNPs, could suggest the presence of multiple mitochondrial haplotypes within an individual. We are unable to test either hypothesis with the available data, because the sequences arise from two different individuals and may be the result of rare or transient structures (Doublet et al., 2013).

Atypical mitochondrial structure, as described here, is patchily distributed across Isopoda (Fig. 1), and sporadically present in various distantly-related aquatic and terrestrial lineages of isopods (Baillie, 2020; Doublet et al., 2012) of both derived and ancestral origin. One hypothesis that has been proposed to explain this distribution, is that atypical structure evolved early in the evolution of Isopoda, prior to the division of sub-orders, and has been subsequently lost via reversion at least three times across the order (Doublet et al., 2012). Our findings provide further support to an early origin of atypical structure, because our research confirms its presence in a marine isopod highly diverged from any other species hitherto known to possess this structure.

The patchy distribution of mitochondrial structures across Isopoda could be explained from a functional perspective by a “terrestrial adaptation hypothesis”. Under this hypothesis, we suggest that atypical structure may have provided an adaptive advantage to the ancestors of modern-day terrestrial isopods in the transition out of a marine environment. Indeed, it has been posited that mitochondrial heteroplasmy introduces additional genetic variation that may play a role in increasing the ability of cells to withstand stressors (Giuliani et al., 2014; Hirose et al., 2018; Leeuwen et al., 2008). Indeed, in the tuatara, mitochondrial duplication has been proposed as a contributor to thermal adaptation, indicating a potential role for structural mitochondrial variants in physiological adaptation (Macey et al., 2021).

The terrestrial adaptation hypothesis is supported by the presence of atypical structure for all terrestrial isopods for which there are data (Fig. 1). A possible exception to this is *Ligia oceanica,* which has contrasting reports of atypical mitochondrial structure (Baillie, 2020; Doublet et al., 2012; Kilpert & Podsiadlowski, 2006). However, this littoral species is arguably semi-aquatic, possesses numerous transitional traits (Michel-Salzat & Bouchon, 2000; Raupach et al., 2014; Schmidt, 2008), and has recently been found to be more closely related to Valvifera or Sphaeromatidea than to the rest of Oniscidea (Dimitriou et al., 2019; Lins et al., 2017). An early origin of atypical structure does not, however, necessarily preclude the hypothesis that atypical structure is adaptive in the terrestrial environment. If this feature did indeed play an adaptive role in the transition to a terrestrial environment, then an early origin and subsequent retention of this feature through selection in the Oniscidea and some Ligiidae, with this trait being lost multiple times among the aquatic isopod suborders, could explain the current distribution of atypical structure. However, it remains to be determined why this trait has been maintained among several aquatic lineages, including *Isocladus*.

An alternative view however, is that the presence of atypical structure across the order is a result of convergent evolution, that is multiple independent origins of atypical structure. This hypothesis has recently gained traction from the recent phylogenetic evidence for multiple independent terrestrial transitions within Isopoda (Dimitriou et al., 2019; Lins et al., 2017). This hypothesis necessitates, however, the repeated duplication of the mitochondrial unit, as well as the evolution of tRNA heteroplasmies (which appear relatively conserved across Isopoda). This explanation is therefore less parsimonious relative to the hypothesis of early origins followed by multiple reversions to typical structure. To distinguish between these hypotheses, it is necessary to determine the mitochondrial arrangement of more species of marine, and transitional marine, species; in particular those within the predominantly supralittoral family Tylidae.

## Conclusion

*Isocladus armatus* possesses an atypical mitochondrial genome consisting of two chromosomes – a 28.7 kb circular dimer, and likely also a 14 kb monomer. The 28.7 kb dimer consists of two copies of the monomer fused together as inverted repeats. Three variant sites occur that differentiate the, otherwise complementary, sides of the 28 kb dimer: the first of these is within a non-coding region, while the other two variants occur in two tRNA genes and cause non-synonymous changes in the tRNA anticodon between sides of the dimer. The presence of this atypical structure in a Sphaeromatid isopod, alongside the differences between sides of the dimer, supports the hypothesis that this structure originated early in the evolution of Isopoda, and the presence of the typical structure in some isopods may be the result of reversion to the ancestral metazoan form.

## Data Accessibility Statement

The assembled mitochondrial unit has been uploaded to GenBank under the accession: OK245257.

## Acknowledgements

We would like to acknowledge the contributions of Stephen Shuster and Christopher Chandler, both of whom provided insight into the mechanisms behind this atypical structure.

## Supplementary Figures

**Supplementary Figure 1:**
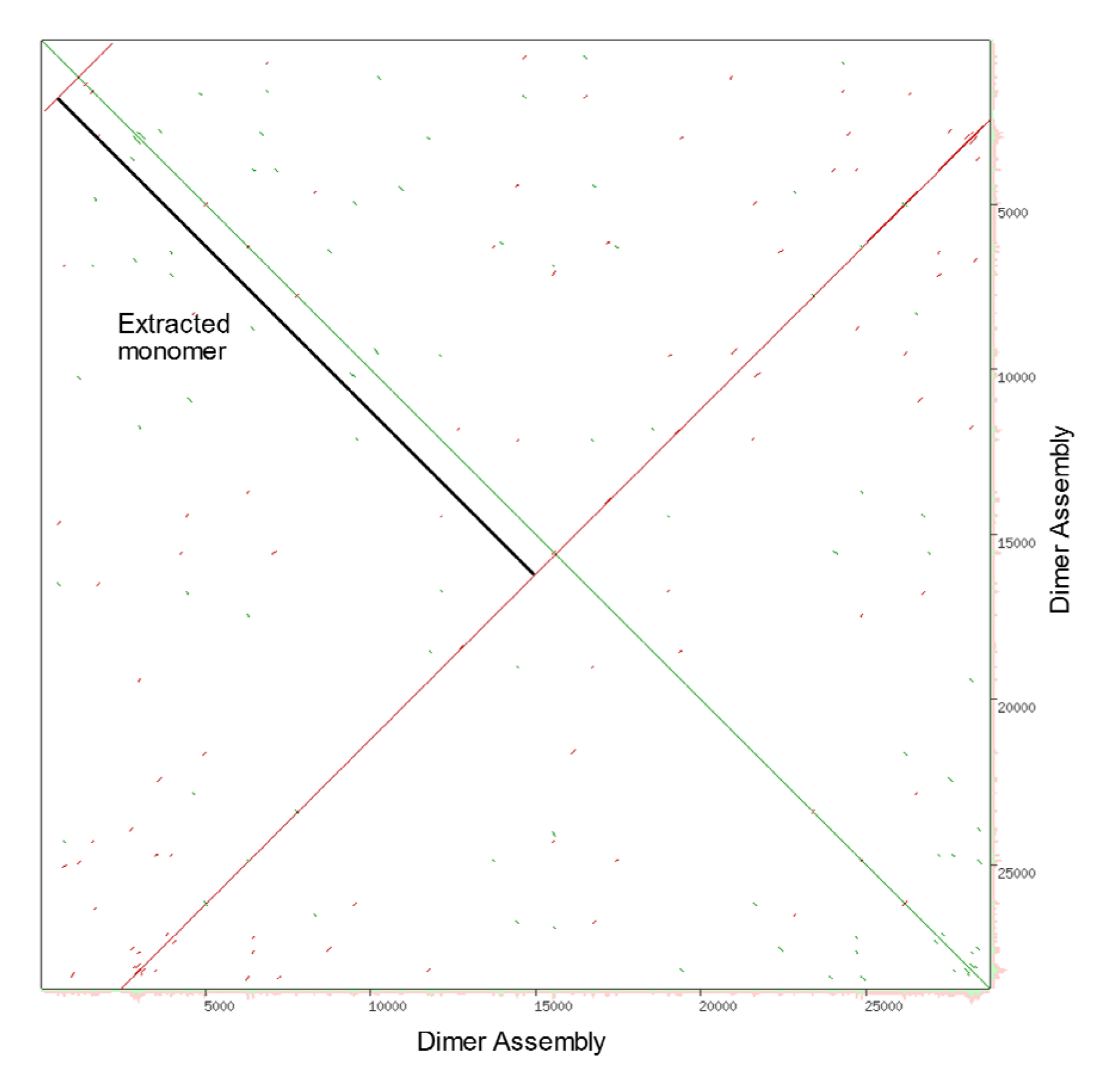
The dotplot used to identify the primary mitochondrial unit, the green line indicates a 1:1 relationship between the forward strand and itself, while the red line indicates the relationship between forward strand and the reverse strand. The black line indicates the section that was extracted and considered the primary mitochondrial unit, inclusive of any unique sequences found within the genome – this is why the red lines do not equal the length of the green line.

**Supplementary Figure 2.**
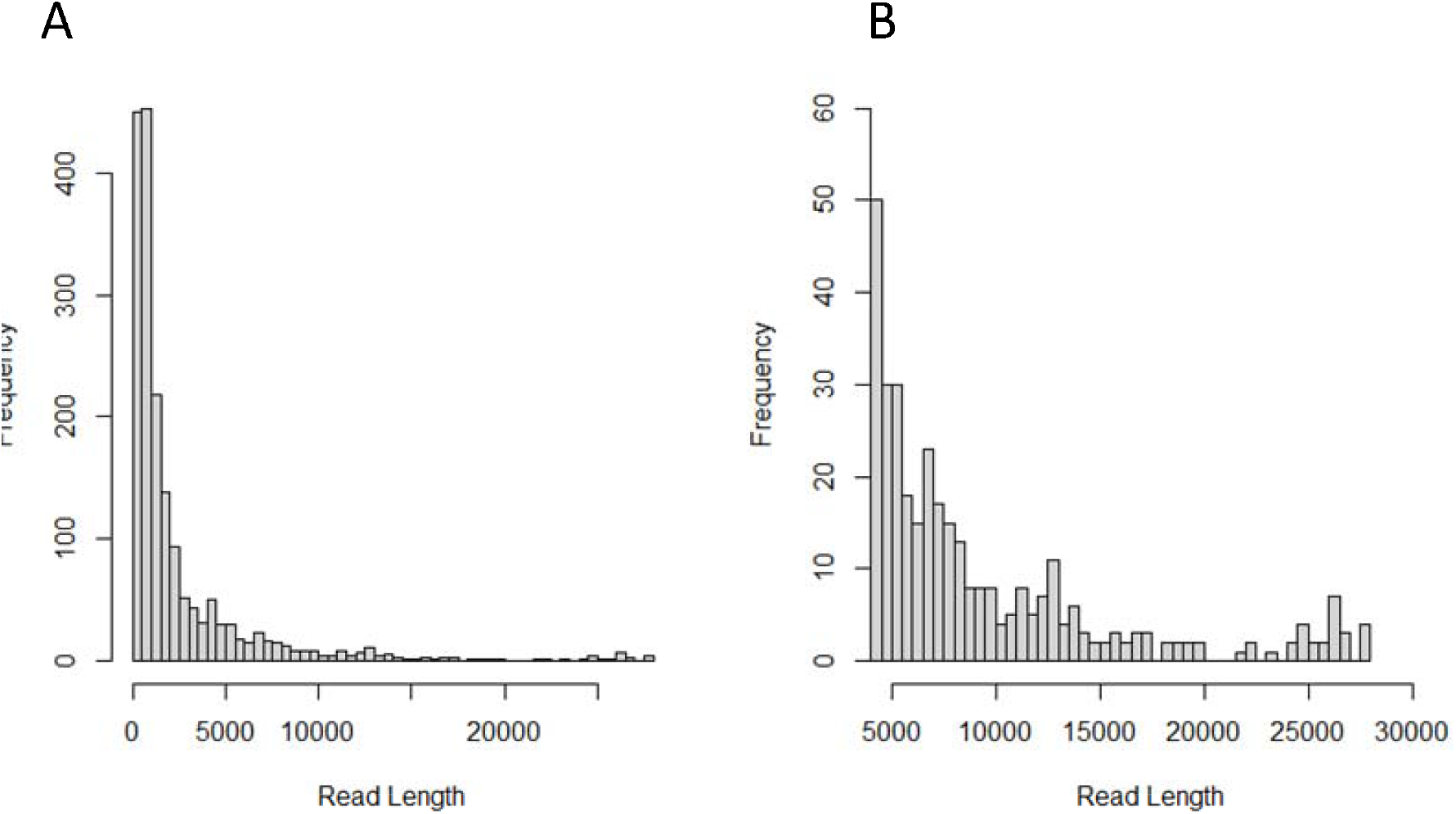
Read length distributions for all mitochondrial originating reads (A) and all mitochondrial originating reads greater than 5000 bp (B).

**Supplementary Figure 3.**
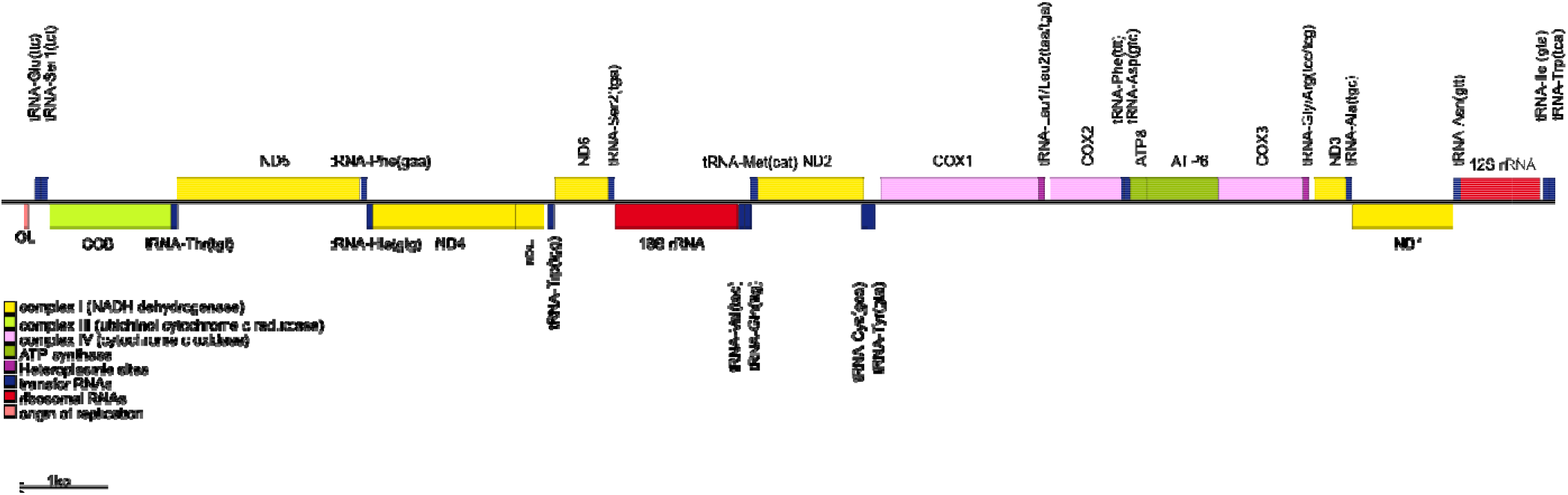
Annotated polished copy of the mitochondrial ‘unit’ for Isocladus armatus. This unit is a total of 14,569 bp, annotations were created using MITOS2, and figure created using OGDRAW2.

## Notes

### Competing Interest Statement

The authors have declared no competing interest.

## References

Al Arab, M., Höner Zu Siederdissen, C., Tout, K., Sahyoun, A. H., Stadler, P. F., & Bernt, M. (2017). Accurate annotation of protein-coding genes in mitochondrial genomes. Molecular Phylogenetics and Evolution, 106, 209–216. https://doi.org/10.1016/j.ympev.2016.09.024

Baillie, C. (2020). Understanding the evolution of cymothoid isopod parasites using comparative genomics and geometric morphometrics [Phd, University of Salford]. http://usir.salford.ac.uk/id/eprint/56173/

Bernt, M., Donath, A., Jühling, F., Externbrink, F., Florentz, C., Fritzsch, G., Pütz, J., Middendorf, M., & Stadler, P. F. (2013). MITOS: improved de novo metazoan mitochondrial genome annotation. Mol. Phylogenet. Evol., 69(2), 313–319. https://doi.org/10.1016/j.ympev.2012.08.023

Burger, G., Gray, M. W., & Franz Lang, B. (2003). Mitochondrial genomes: Anything goes. Trends in Genetics, 19(12), 709–716. https://doi.org/10.1016/j.tig.2003.10.012

Chandler, C. H., Badawi, M., Moumen, B., Grève, P., & Cordaux, R. (2015). Multiple Conserved Heteroplasmic Sites in tRNA Genes in the Mitochondrial Genomes of Terrestrial Isopods (Oniscidea). G3, 5(7), 1317–1322.

Dimitriou, A. C., Taiti, S., & Sfenthourakis, S. (2019). Genetic evidence against monophyly of Oniscidea implies a need to revise scenarios for the origin of terrestrial isopods. Scientific Reports, 9(1), 18508. https://doi.org/10.1038/s41598-019-55071-4

Doublet, V., Helleu, Q., Raimond, R., Souty-Grosset, C., & Marcadé, I. (2013). Inverted repeats and genome architecture conversions of terrestrial isopods mitochondrial DNA. J. Mol. Evol., 77(3), 107–118.

Doublet, V., Raimond, R., Grandjean, F., Lafitte, A., Souty-Grosset, C., & Marcadé, I. (2012). Widespread atypical mitochondrial DNA structure in isopods (Crustacea, Peracarida) related to a constitutive heteroplasmy in terrestrial species. Genome, 55(3), 234–244.

Doublet, V., Ubrig, E., Alioua, A., Bouchon, D., Marcadé, I., & Maréchal-Drouard, L. (2015). Large gene overlaps and tRNA processing in the compact mitochondrial genome of the crustacean Armadillidium vulgare. RNA Biology, 12(10), 1159–1168. https://doi.org/10.1080/15476286.2015.1090078

Drummond, A. J., Ashton, B., Buxton, S., Cheung, M., Cooper, A., Duran, C., Field, M., Heled, J., Kearse, M., Markowitz, S., & Others. (2011). Geneious. Biomatters Ltd.

Giuliani, C., Barbieri, C., Li, M., Bucci, L., Monti, D., Passarino, G., Luiselli, D., Franceschi, C., Stoneking, M., & Garagnani, P. (2014). Transmission from centenarians to their offspring of mtDNA heteroplasmy revealed by ultra-deep sequencing. Aging, 6(6), 454–467. https://doi.org/10.18632/aging.100661

Hirose, M., Schilf, P., Gupta, Y., Zarse, K., Künstner, A., Fähnrich, A., Busch, H., Yin, J., Wright, M. N., Ziegler, A., Vallier, M., Belheouane, M., Baines, J. F., Tautz, D., Johann, K., Oelkrug, R., Mittag, J., Lehnert, H., Othman, A., … Ibrahim, S. M. (2018). Low-level mitochondrial heteroplasmy modulates DNA replication, glucose metabolism and lifespan in mice. Scientific Reports, 8(1), 5872. https://doi.org/10.1038/s41598-018-24290-6

Kilpert, F., Held, C., & Podsiadlowski, L. (2012). Multiple rearrangements in mitochondrial genomes of Isopoda and phylogenetic implications. Mol. Phylogenet. Evol., 64(1), 106–117.

Kilpert, F., & Podsiadlowski, L. (2006). The complete mitochondrial genome of the common sea slater, Ligia oceanica (Crustacea, Isopoda) bears a novel gene order and unusual control region features. BMC Genomics, 7(1), 241. https://doi.org/10.1186/1471-2164-7-241

Kolmogorov, M., Yuan, J., Lin, Y., & Pevzner, P. A. (2019). Assembly of long, error-prone reads using repeat graphs. Nat. Biotechnol., 37(5), 540–546.

Lavrov, D. V., & Pett, W. (2016). Animal Mitochondrial DNA as We Do Not Know It: Mt-Genome Organization and Evolution in Nonbilaterian Lineages. Genome Biology and Evolution, 8(9), 2896–2913. https://doi.org/10.1093/gbe/evw195

Leeuwen, T. V., Vanholme, B., Pottelberge, S. V., Nieuwenhuyse, P. V., Nauen, R., Tirry, L., & Denholm, I. (2008). Mitochondrial heteroplasmy and the evolution of insecticide resistance: Non-Mendelian inheritance in action. Proceedings of the National Academy of Sciences, 105(16), 5980–5985. https://doi.org/10.1073/pnas.0802224105

Li, H. (2011). A statistical framework for SNP calling, mutation discovery, association mapping and population genetical parameter estimation from sequencing data. Bioinformatics (Oxford, England), 27(21), 2987–2993. https://doi.org/10.1093/bioinformatics/btr509

Li, H., & Durbin, R. (2009). Fast and accurate short read alignment with Burrows–Wheeler transform. Bioinformatics, 25(14), 1754–1760. https://doi.org/10.1093/bioinformatics/btp324

Lins, L. S. F., Ho, S. Y. W., & Lo, N. (2017). An evolutionary timescale for terrestrial isopods and a lack of molecular support for the monophyly of Oniscidea (Crustacea: Isopoda). Organisms Diversity & Evolution, 17(4), 813–820. https://doi.org/10.1007/s13127-017-0346-2

Lins, L. S. F., Ho, S. Y. W., Wilson, G. D. F., & Lo, N. (2012). Evidence for Permo-Triassic colonization of the deep sea by isopods. Biol. Lett., 8(6), 979–982.

Liu, H., Li, H., Song, F., Gu, W., Feng, J., Cai, W., & Shao, R. (2017). Novel insights into mitochondrial gene rearrangement in thrips (Insecta: Thysanoptera) from the grass thrips, Anaphothrips obscurus. Sci. Rep., 7(1), 4284.

Macey, J. R., Pabinger, S., Barbieri, C. G., Buring, E. S., Gonzalez, V. L., Mulcahy, D. G., DeMeo, D. P., Urban, L., Hime, P. M., Prost, S., Elliott, A. N., & Gemmell, N. J. (2021). Evidence of two deeply divergent co-existing mitochondrial genomes in the Tuatara reveals an extremely complex genomic organization. Communications Biology, 4(1), 1–10. https://doi.org/10.1038/s42003-020-01639-0

Michel-Salzat, A., & Bouchon, D. (2000). Phylogenetic analysis of mitochondrial LSU rRNA in oniscids. Comptes Rendus de l’Académie Des Sciences - Series III - Sciences de La Vie, 323(9), 827–837. https://doi.org/10.1016/S0764-4469(00)01221-X

Noé, L., & Kucherov, G. (2005). YASS: Enhancing the sensitivity of DNA similarity search. Nucleic Acids Research, 33(Web Server issue), W540–W543. https://doi.org/10.1093/nar/gki478

Pearman, W. S., Wells, S. J., Silander, O. K., Freed, N. E., & Dale, J. (2020). Concordant geographic and genetic structure revealed by genotyping-by-sequencing in a New Zealand marine isopod. Ecology and Evolution, 10(24), 13624–13639. https://doi.org/10.1002/ece3.6802

Peccoud, J., Chebbi, M. A., Cormier, A., Moumen, B., Gilbert, C., Marcadé, I., Chandler, C., & Cordaux, R. (2017). Untangling Heteroplasmy, Structure, and Evolution of an Atypical Mitochondrial Genome by PacBio Sequencing. Genetics, 207(1), 269–280.

Raimond, R., Marcadé, I., Bouchon, D., Rigaud, T., Bossy, J.-P., & Souty-Grosset, C. (1999). Organization of the Large Mitochondrial Genome in the Isopod Armadillidium vulgare. Genetics, 151(1), 203–210.

Raupach, M. J., Bininda-Emonds, O. R. P., Knebelsberger, T., Laakmann, S., Pfaender, J., & Leese, F. (2014). Phylogeographical analysis of Ligia oceanica (Crustacea: Isopoda) reveals two deeply divergent mitochondrial lineages. Biological Journal of the Linnean Society, 112(1), 16–30. https://doi.org/10.1111/bij.12254

Robinson, J. T., Thorvaldsdóttir, H., Winckler, W., Guttman, M., Lander, E. S., Getz, G., & Mesirov, J. P. (2011). Integrative genomics viewer. Nature Biotechnology, 29(1), 24–26. https://doi.org/10.1038/nbt.1754

Sahyoun, A. H., Hölzer, M., Jühling, F., Höner zu Siederdissen, C., Al-Arab, M., Tout, K., Marz, M., Middendorf, M., Stadler, P. F., & Bernt, M. (2015). Towards a comprehensive picture of alloacceptor tRNA remolding in metazoan mitochondrial genomes. Nucleic Acids Research, 43(16), 8044–8056. https://doi.org/10.1093/nar/gkv746

Schmidt, C. (2008). Phylogeny of the Terrestrial Isopoda (Oniscidea): A Review. Arthropod Systematics & Phylogeny, 66(2), 191–226.

Vaser, R., Soviċ, I., Nagarajan, N., & Šikić, M. (2017). Fast and accurate de novo genome assembly from long uncorrected reads. Genome Research, 27(5), 737–746. https://doi.org/10.1101/gr.214270.116

Wei, D.-D., Shao, R., Yuan, M.-L., Dou, W., Barker, S. C., & Wang, J.-J. (2012). The multipartite mitochondrial genome of Liposcelis bostrychophila: Insights into the evolution of mitochondrial genomes in bilateral animals. PLoS One, 7(3), e33973.

Yang, M., Gao, T., Hui, D., Chen, X., & Liu, W. (2019). Complete mitochondrial genome and the phylogenetic position of the Sphaeroma sp. (Crustacea, Isopod, Sphaeromatidae). Mitochondrial DNA Part B, 4(2), 3896–3897. https://doi.org/10.1080/23802359.2019.1687350

Yang, M., Gao, T., Yan, B., Chen, X., & Liu, W. (2019). Complete mitochondrial genome and the phylogenetic position of a wood-boring Isopod Sphaeroma terebrans (Crustacea, Isopod, Sphaeromatidae). Mitochondrial DNA Part B, 4(1), 1920–1921.

Zou, H., Jakovliċ, I., Zhang, D., Chen, R., Mahboob, S., Al-Ghanim, K. A., Al-Misned, F., Li, W.-X., & Wang, G.-T. (2018). The complete mitochondrial genome of Cymothoa indica has a highly rearranged gene order and clusters at the very base of the Isopoda clade. PLoS One, 13(9), e0203089.

